# Global ecotypes in the ubiquitous marine clade SAR86

**DOI:** 10.1101/635185

**Authors:** Adrienne Hoarfrost, Stephen Nayfach, Joshua Ladau, Shibu Yooseph, Carol Arnosti, Chris L Dupont, Katherine S. Pollard

## Abstract

SAR86 is an abundant and ubiquitous heterotroph in the surface ocean that plays a central role in the function of marine ecosystems. We hypothesized that despite its ubiquity, different SAR86 subgroups may be endemic to specific ocean regions and functionally specialized for unique marine environments. However, the global biogeographical distributions of SAR86 genes, and the manner in which these distributions correlate with marine environments, have not been investigated. We quantified SAR86 gene content across globally-distributed metagenomic samples and modeled these gene distributions as a function of 51 environmental variables. We identified five distinct clusters of genes within the SAR86 pangenome, each with a unique geographic distribution associated with specific environmental characteristics. Gene clusters are characterized by strong taxonomic enrichment of distinct SAR86 genomes and partial assemblies, as well as differential enrichment of certain functional groups, suggesting differing functional and ecological roles of SAR86 ecotypes. We then leveraged our models and high-resolution, remote sensing-derived environmental data to predict the distributions of SAR86 gene clusters across the world’s oceans, creating global maps of SAR86 ecotype distributions. Our results reveal that SAR86 exhibits previously unknown, complex biogeography, and provide a framework for exploring geographic distributions of genetic diversity from other microbial clades.

## Introduction

Marine microbes are important drivers of biogeochemical cycling and ecological function [1, 2]. Many studies have demonstrated the link between microbial genetic diversity and functional capacities [e.g. [3–7], as well as the dependence of microbial community structure and function on environmental variables [5, 8, 9]. However, the complexity of microbial communities and of their interactions with their environment limit our ability to link microbial genetic and functional variation across environments [10]. Furthermore, we have only limited understanding of the geographic distributions of genetic diversity within key taxa, the relationship of gene distributions to environmental conditions, and the manner in which these distributions may result in distinct ecotypes across different environments and regions. Our limitations in mapping microbial genetic diversity to geographic distributions restrict our ability to predict microbial ecotypes across the environment. Accurate models linking environmental and microbial variables may improve our current ability to incorporate biological inputs into ecosystem models, which often rely on simplified biological systems utilizing incomplete environmental relationships or imprecise evaluations of the functional capabilities of microbial communities at different locations [11, 12].

In microbial ecology, an ecotype [13] is often identified in practice as a group of closely related lineages that co-occur on the same spatial or temporal scale and are associated with particular environmental conditions. This contrasts with the classical ecological definition, which additionally specifies that an ecotype must be genotypically adapted to the environmental conditions it is associated with [14]. In microbial ecology, where community members often lack cultured representatives and experiments directly measuring adaptive capacity to manipulated environmental conditions are challenging to conduct, adaptation is often difficult to demonstrate conclusively. In this study, we define an ecotype to be a group of lineages within a clade whose genomes contain a similar set of genes with a common geographic distribution associated with distinct environmental conditions. This definition is consistent with previous studies of microbial ecotypes [15]. Additionally, we require an ecotype to be taxonomically and functionally differentiated from other ecotypes, which may indicate an adaptive strategy specific to that ecotype, although we do not explicitly test for genetic signatures of adaptation.

The biogeography of marine microbes has been observed at scales from single depth profiles [4] to global surveys [16, 17], revealing spatial and temporal patterns in microbial community structure [16, 18], function [8, 19], and diversity [17]. Many marine microbial clades exhibit population structure that correlates with their differential geographic distributions [20]. Because most microbes have large pangenomes and flexible gene content [20], there is significant interest in elucidating the differential functional capabilities of microbial ecotypes and mapping their biogeographical distributions. Associating geographic distributions of microbial ecotypes with environmental conditions could illuminate the links between microbial community structure, function, and ecosystem processes, enabling predictions of biological and chemical shifts in the world’s oceans as environmental conditions change. However, there have been very few efforts to predict biogeographic patterns of genetic and functional diversity of key microbial taxa at large spatial scales in the ocean [17, 21].

SAR86 is a ubiquitous marine heterotroph frequently found in surface waters, classified by their 16S rRNA gene similarity as a clade within the Gammaproteobacteria [22–24]. SAR86 is a very diverse group with at least three subclades [23, 24]. Despite its ubiquity in marine systems, SAR86 eludes cultivation, and therefore knowledge of the ecological role of SAR86 in marine microbial communities is limited to evidence from genomes curated from single-cell sequencing or metagenomic assembly [25–27]. These genomes suggest that SAR86 gene sets, and hence functional capabilities, vary greatly across locations, even though the clade is very commonly detected in marine environments. However, little is known about the manner in which the distribution of subspecies and the vast genetic diversity within the SAR86 pangenome may vary across large spatial extents, and what environmental factors may affect the geographic distributions of different SAR86 gene families.

In this study, we build a custom pangenome of SAR86 genes from metagenomic co-assemblies and five available reference genomes. We then quantify the presence of each gene in the pangenome across diverse marine epipelagic waters using hundreds of publicly available, globally-distributed shotgun metagenomes. We find that geographic distributions of SAR86 genes are strongly associated with environmental variables, and we leverage these associations to build machine learning models that accurately predict the presence of SAR86 genes from environmental data. Using global-scale environmental measurements from satellite and shipboard sources, we use our models to predict the global distribution of each geographically variable gene in the SAR86 pangenome at a 9km^2^ resolution. Our machine learning approach enables patterns in the environmental variables that best predict the distributions of SAR86 genes to emerge from the global metagenomic dataset without explicitly assuming *a priori* relationships between inputs and outputs. Analysis of the resultant models reveals five clusters of genes with unique environmental and geographic distributions, defining five ecotypes within the SAR86 clade. We conclude that patterns of taxonomic and functional enrichment across these ecotypes reveal previously underappreciated complexity in the geographic distributions underlying the pangenome of this otherwise ubiquitous marine heterotroph, with great potential to illuminate structure-function relationships across the marine environment.

## Materials & Methods

### Creation of the SAR86 pangenome and global SAR86 gene presence/absence dataset

A custom pangenome of 51 711 nonredundant SAR86 genes was created with the MIDAS tool [20], from a combination of genomic sources [23, 24, 25] as well as a massive co-assembly of metagenomic sequences (Supplemental Text 1.1-1.2).

A global dataset of SAR86 gene presence/absence for each gene in the SAR86 pangenome was then created. Shotgun metagenomic sequencing reads from the TARA project [9] were mapped to the SAR86 pangenome, and the resulting normalized read coverage for each gene was used to determine SAR86 gene presence or absence for all SAR86 genes at 198 TARA sites (Supplemental Text 1.3).

### Environmental data curation and processing

In order to build models predicting SAR86 gene presence from environmental variables, environmental data available at resolution between 9km to 1-degree and at global scale were curated from a combination of contemporary satellite data and historical averages of satellite and interpolated in situ measurements. A total of 51 environmental features were compiled (SI Table 1, Supplemental Text 1.4). Normalized environmental feature values closest to each TARA site’s latitude, longitude, and, where relevant, sampling depth and/or sampling date (SI Table 2) served as the input feature vectors for each TARA site during model training.

### Gene presence/absence models & predictions

Classification models predicting SAR86 gene presence or absence as a function of the environmental feature vectors across TARA sites were built for each of 24 317 geographically variable SAR86 genes, using logistic regression with L1 regularization (Supplemental Text 1.5). Geographically variable genes were defined as genes present at between 20-80% of TARA sites. 155 TARA sites for which SAR86 was present and environmental data was available were split into training, validation, and test sets of 111, 13, and 31 sites respectively. The final models trained independently for each of the 24 317 geographically variable genes can be reproduced with code available on the associated Github repository [29].

### Clustering, global maps of ecotypes, & enrichment analysis

To identify groups of SAR86 genes whose geographic distributions are best predicted by similar environmental variables, we clustered genes into 5 clusters on the logistic regression model coefficients for each environmental feature using a k-means algorithm (Supplemental Text 1.6). Clustering on environmental features associated with gene models enabled us to identify the environmental variables underlying geographic distributions of genes, and also enabled the projection of predicted cluster distributions at global scales. To produce global projections (i.e., maps) of each SAR86 gene cluster, we predicted the presence or absence of each cluster at 9km2 resolution and global scale from the available satellite and historical environmental data ([29], Supplemental Text 1.6). A Jupyter notebook and a python script for reproducing clusters and cluster projections are available ([29]).

The distribution and enrichment across clusters were evaluated at the genome, contig, and functional level for two SAR86 reference genomes SAR86A and SAR86E, for the contigs of the SAR86 co-assembly, and for the functional annotations to Pfam [30] for the SAR86 pangenome (Supplemental Text 1.7). This produced a vector of taxonomic/functional enrichment values associated with each contig/annotation for each cluster, with which the statistical significance of cluster enrichment could be tested (Supplemental Text 1.7).

## Results

This study first modeled the relationships between SAR86 gene distributions and environmental variables. We used a regularized logistic regression approach to identify the subset of environmental variables that are most important for predicting the geographical distributions of each gene and to estimate the strength of these gene-environmental variable relationships. Using unsupervised clustering of these association profiles, we then identified clusters of genes with similar environmental distributions. Clustering enabled us to identify the structure underlying the environmental gene distributions without explicit prior knowledge of expected SAR86 ecotypes. By using environmental variables available at global scale, we leveraged our gene models to predict the geographic distribution of these emergent ecotypes in regions far beyond the sampling locations specific to the TARA study.

### Accurate prediction of SAR86 gene distributions from environmental variables

SAR86 gene content in TARA Oceans metagenomes is associated with environmental characteristics of the sampling locations. We built a regularized logistic regression model for each gene that accurately predicts the probability of the gene being present at a given location as a function of the most predictive subset of environmental variables (Methods, Supplemental Text 1.5).

The resulting 24 317 gene models predict SAR86 gene presence/absence with an average of 79.4% accuracy in the test set, and a median test accuracy of 80.6%. Precision and recall measures are roughly even (0.85 and 0.81, respectively; SI Fig 3a), with an F1 score of 0.83. For 21 264 out of 24 317 genes (87.4%), the models have accuracies in the test set that are an improvement over the majority class accuracy – the accuracy of the model if it predicts ‘always absent’ or ‘always present’, whichever is in the majority (SI Fig 3b).

As an additional test of the robustness of the models, the accuracy of predictions at those TARA sites that were not included in model development, where SAR86 was not present or were in very low abundance, was also examined. There were 20 such sites for which environmental data was available for all features. These 20 sites were primarily mesopelagic samples, distributed across all ocean basins (Supplemental Text 1.5). Across these 20 sites, the average accuracy of the gene models is 68.5%, while the median accuracy is 70.0%. While this performance is below that achieved at sites where SAR86 was present, it suggests that our models are able to make fairly accurate predictions even when extrapolating outside of the distribution of gene presence used in training.

An average of 17 of 51 environmental features is significantly associated with each gene’s distribution across TARA Oceans sites. Across multiple gene models, the same environmental feature was frequently selected during model training (SI Fig 4). These frequently associated variables include latitude, longitude, distance from land, ocean depth, and other features that might describe the general ocean basin or region of a sample; as well as pH, sea surface temperature, pycnocline depth, nitrogen:phosphorous ratio, cloud fraction, and other environmental factors that describe regions of the ocean that experience particular environmental conditions.

While the environmental features that best predict gene presence/absence vary by the individual gene model, and many of the 51 environmental variables covary with one another, training logistic regression multiple times on the same data with different random seeds resulted in the same sets of environmental features being chosen as the most predictive for each gene model (see Jupyter notebook in [29]). This consistency suggests that the environmental features selected in each model reflect a true difference in predictive power between the selected features and those that were not selected, rather than a random choice among features that are roughly equally predictive.

### Clustering of SAR86 genes into common environmental distributions & global projections of their biogeographic distributions

The environmental features that best predict individual genes, and the strength of the coefficients associated with any particular environmental feature, vary by the individual gene model. However, there are apparent patterns among genes, with some groups of genes appearing to be predicted by similar environmental variables, as well as similar magnitudes and signs of the coefficients associated with those variables. These patterns suggest that genes that are predicted by similar environmental features occupy similar geographic distributions characterized by unique environmental conditions.

K-means clustering of genes by their logistic regression environmental feature coefficients identified five clusters within the SAR86 pangenome characterized by similar environmental distributions (Fig 1). The average environmental feature coefficient across all genes in each cluster (the “centroid”) demonstrates the distinct pattern of association with environmental features of each cluster (SI Table 3).

**Fig. 1.**
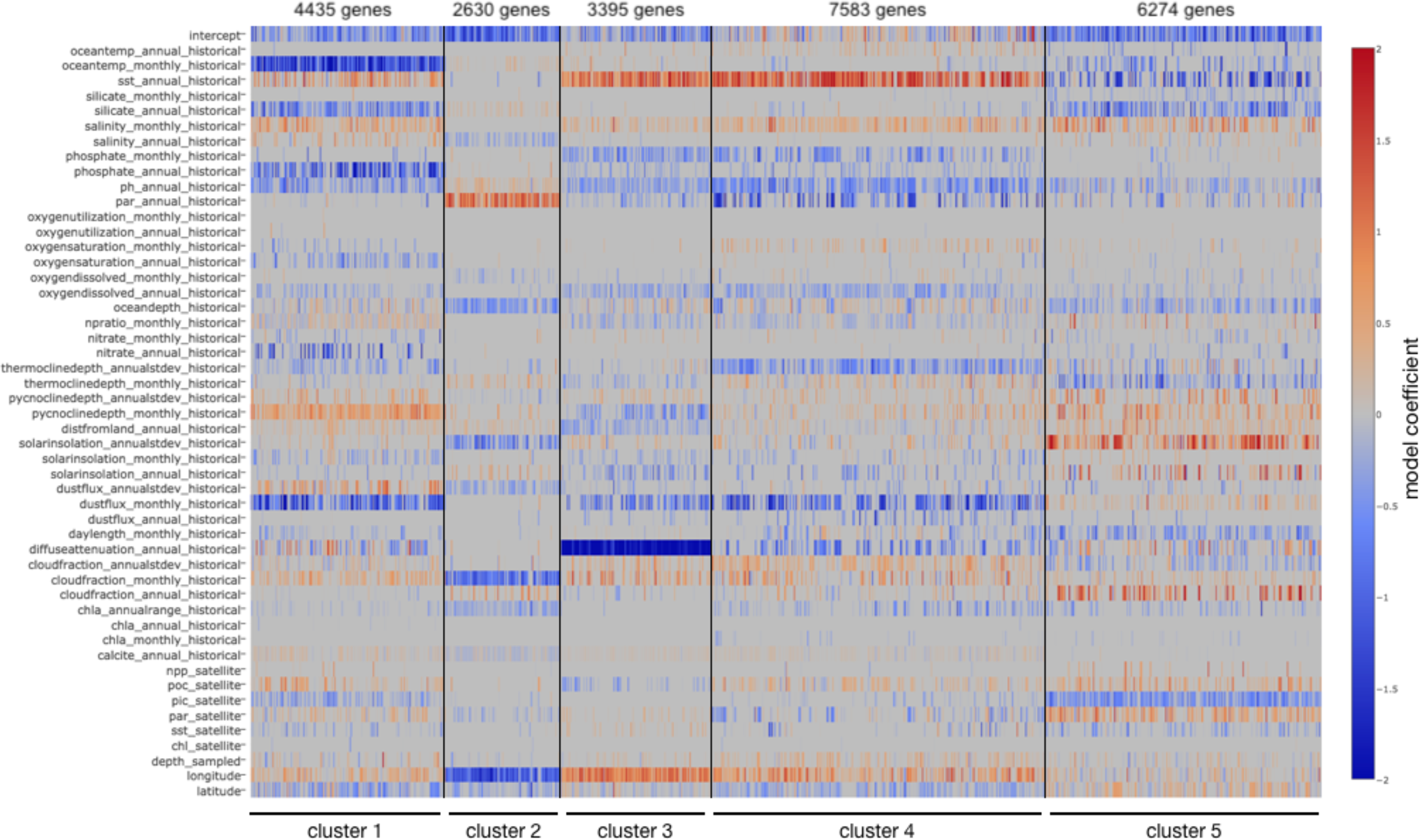
Heatmap of model coefficients for each environmental feature (rows) and gene (columns), ordered by cluster (x axis).

Each TARA site contains genes from a mixture of clusters, but the dominant clusters and the evenness of the proportion of each cluster is variable across sites (Fig 2, SI Fig 5, SI Table 4). For example, cluster 2 is strongly associated with longitudes in the western hemisphere, and this is also reflected across TARA samples, for which cluster 2 is present in highest proportions for those TARA sites sampled in the Pacific Ocean (Fig 2, SI Fig 5b). In contrast, cluster 3 genes are found in higher proportions at TARA sites sampled in the eastern hemisphere, reflecting their predicted geographic distributions (Fig 2, SI Fig 5c).

**Fig. 2.**
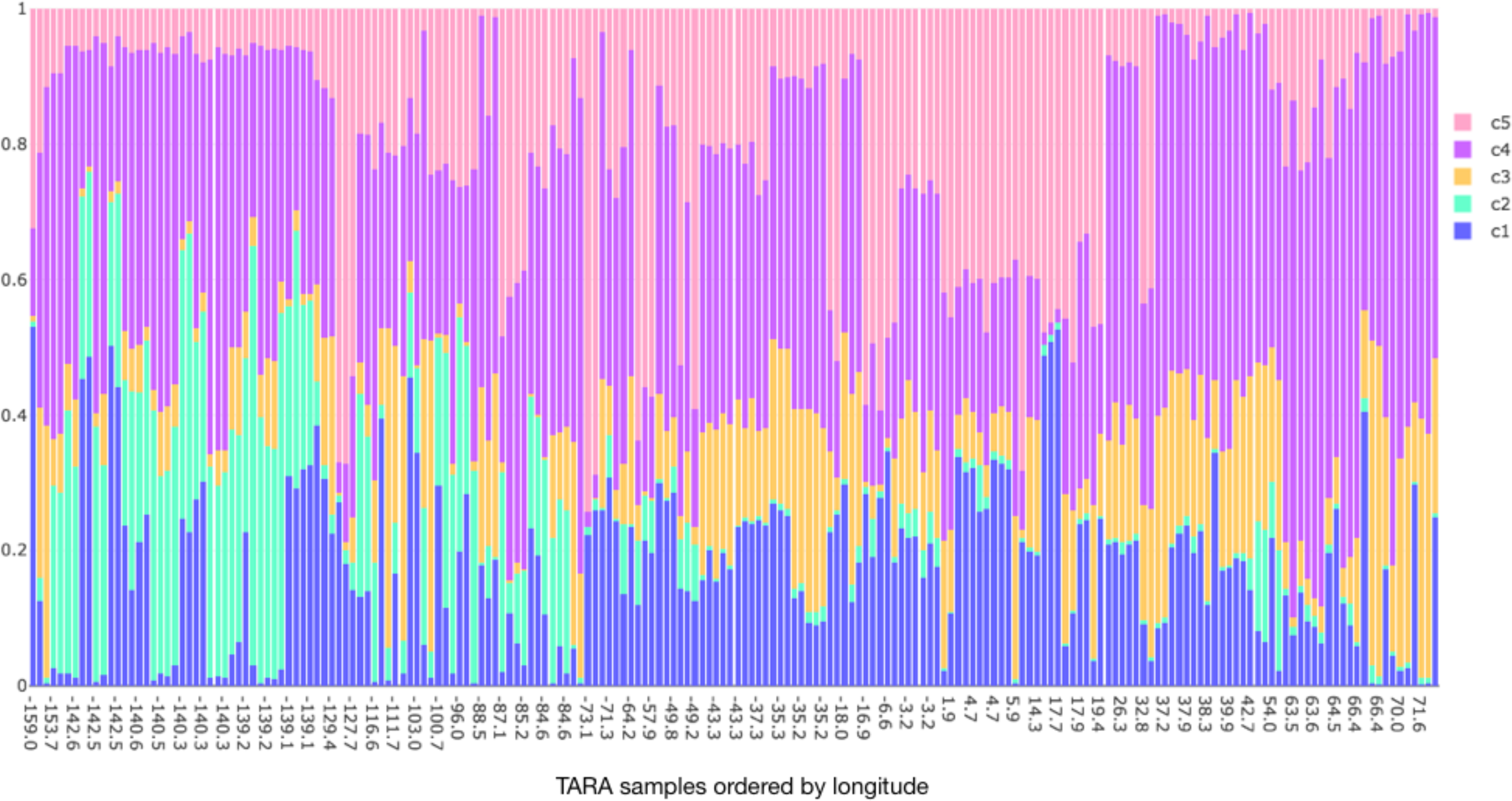
Relative proportion of clusters at each TARA site (vertical bars). TARA sites are sorted by longitude (x axis; negative numbers correspond to longitude west of the prime meridian). Blue, cluster 1; green, cluster 2; yellow, cluster 3; purple, cluster 4; pink, cluster 5.

A Shannon diversity metric was used to measure the relative evenness and proportion of the five clusters at each TARA site (SI Table 4, Supplemental Text 1.7). The TARA sites with the lowest Shannon diversity include TARA station 93 at 34°S and 73°W off the coast of Chile, which is dominated by cluster 5 genes, and TARA stations 38, 42, 45, and 36 in the Indian Ocean, which are dominated by cluster 4 genes. The TARA sites with the highest Shannon diversity include many of the mesopelagic depth samples in the Pacific Ocean, as well as station 70 in the South Atlantic basin at 20.4°S and 3.2°W.

We next used the cluster centroids and global-scale environmental data to predict the geographic distribution of each cluster beyond the TARA sampling locations (Fig 3). These global projections reveal the differential distributions of SAR86 gene clusters. These differential distributions are reflected in variation across longitude (e.g. cluster 2 versus clusters 3 and 4), latitude (e.g. clusters 1 and 5 versus clusters 2, 3, and 4), and season (e.g. cluster 1, Fig 3). In each case, the highest magnitude coefficients for each cluster are suggestive of their predicted geographic distributions (SI Table 3, Supplemental Text 2.1).

**Fig. 3.**
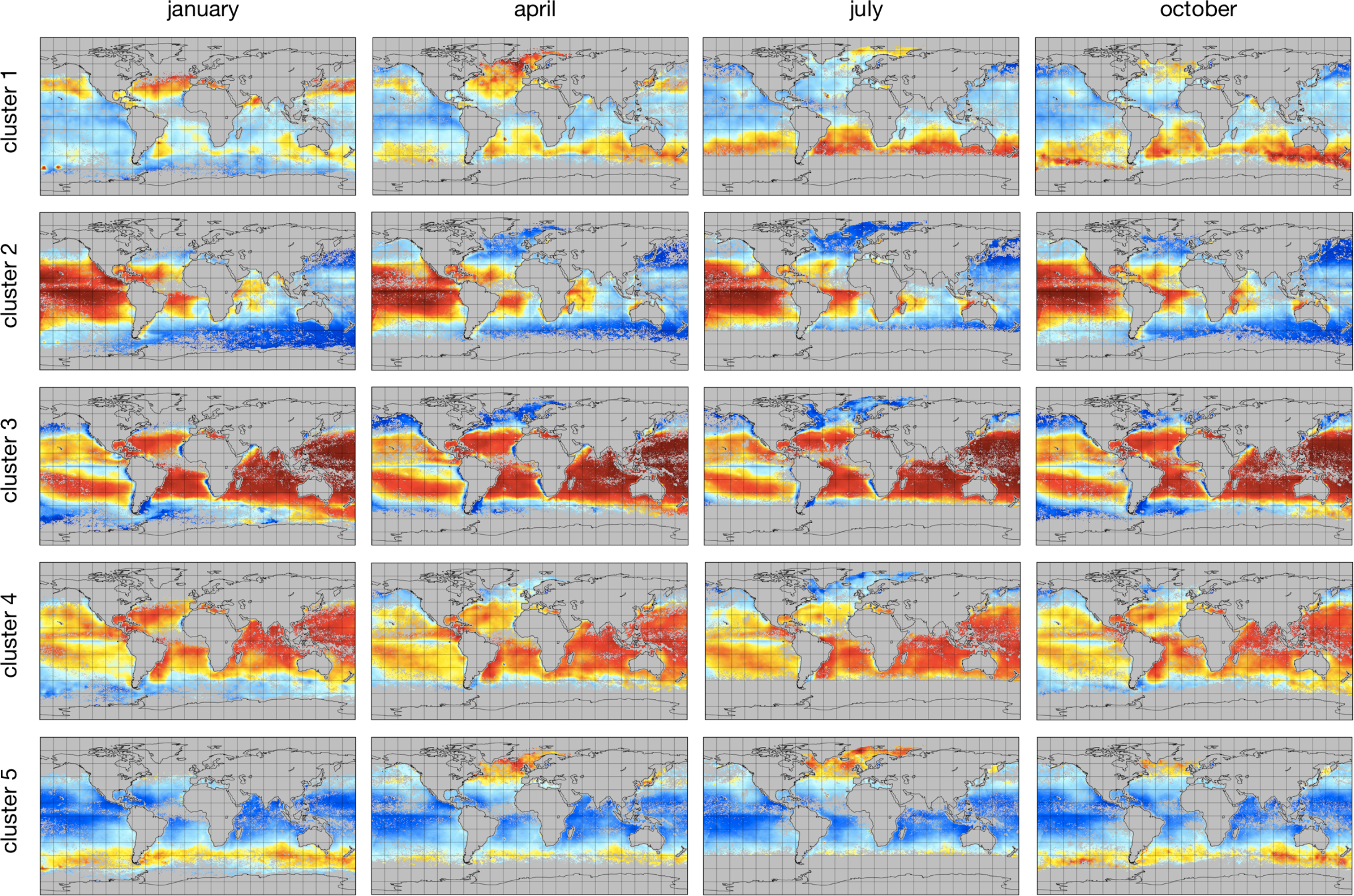
Global predictions of SAR86 gene cluster distributions for each cluster (rows) in January, April, July, and October of 2009 (columns). Red indicates a high confidence of a gene cluster being present, blue a high confidence of a gene cluster being absent, and white a low confidence prediction.

### Taxonomic enrichment & functional differentiation across clusters define SAR86 ecotypes

The cluster assignments of genes from the SAR86 reference genomes SAR86A and SAR86E show clear partitioning on taxonomic lines. Genes from each genome are assigned primarily to two clusters, and each cluster is dominated by one genome. SAR86A genes are partitioned primarily into clusters 4 and 3, with 493 and 118 out of the 622 SAR86A genes assigned to cluster 4 and 3 respectively, while only 4 and 7 genes were assigned to clusters 2 and 5, and 0 genes to cluster 1. The 157 SAR86E genes were partitioned into clusters 1 and 5, with 76 and 78 genes respectively, while only 2 and 1 genes were assigned to clusters 2 and 4, respectively, and 0 genes to cluster 3.

Clusters also show clear taxonomic differentiation at the contig level. Those genes that do not originate from one of the five SAR86 genomes constitute a total length of 22 Mbp originating from 732 contigs from the SAR86 co-assembly. All clusters are significantly enriched in specific contigs (p<0.001, Fig 4c), with a unique set of contigs enriched on each cluster. Genes from the same contig are generally assigned to the same cluster, such that gene assignments of almost all contigs, 540 out of 732 contigs, are enriched on only one cluster, 183 contigs are enriched on only two clusters, and the remaining 9 contigs are enriched on 3 clusters (Fig 4). Where a contig is enriched, the enrichment is strong, with an average enrichment of 3.03 and a standard deviation of 0.43, and ranging from 1.41 in cluster 4 to 5.25 in cluster 2.

**Fig. 4.**
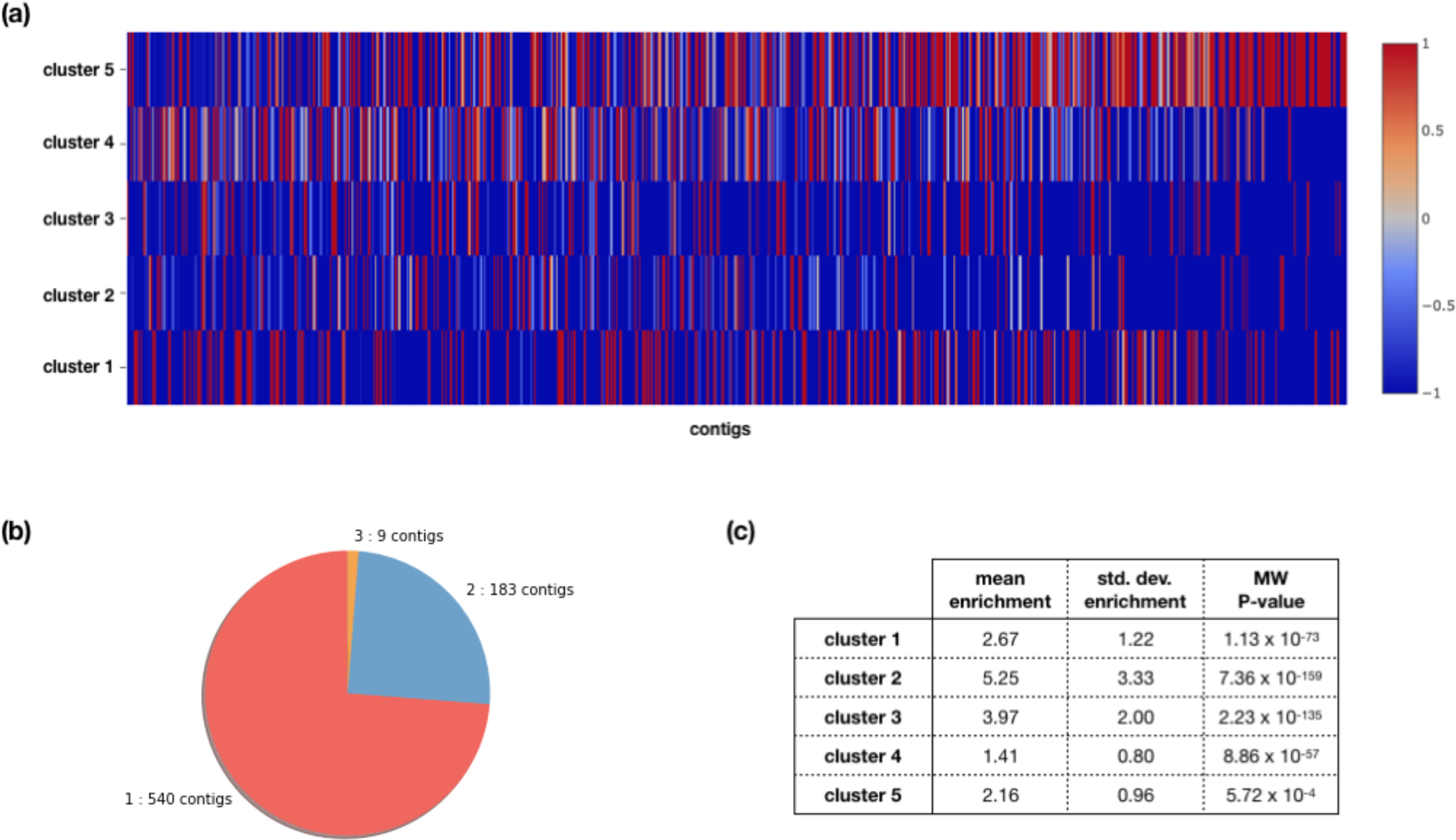
Contig enrichment in clusters. (a) Heatmap of enrichment (red) or depletion (blue) of each contig (columns) across each cluster (rows). (b) Pie chart of the number of clusters in which SAR86 contigs are enriched. (c) Mean positive enrichment value, standard deviation of positive enrichment values, and the Mann-Whitney P value for significance of cluster enrichment, for each cluster.

The taxonomic partitioning of clusters is also evident in their distribution across TARA sites. First, the cluster proportions and the relative abundances of SAR86 genomes at TARA sites reflect the taxonomic differentiation of genomes across clusters. The clusters associated with SAR86A (clusters 3 and 4) are in higher proportions relative to the clusters associated with SAR86E (clusters 1 and 5) at TARA sites where SAR86A abundances are higher relative to SAR86E (SI Fig 6, Pearson R2 = 0.70, P=1.56×10-26). In addition to this genomic evidence, the normalized read coverage across TARA sites for genes from the same cluster are more highly correlated with one another than genes from different clusters (SI Fig 7), as would be expected if genes belonging to the same cluster share a common taxonomic origin. This indicates that genes from the same genome are assigned to the same cluster, although a single cluster may be made up of genes from multiple genomes. Indeed, the 22Mbp of genomic material in the SAR86 co-assembly is enough for at least 11 genomes of size similar to that of known SAR86 reference genomes, so multiple genomes are expected to be contained within the 5 identified clusters. These clusters are thus composed of genes that co-occur with one another across similar environmental contexts, and are taxonomically differentiated, but do not necessarily represent individual SAR86 genomes.

In addition to taxonomic enrichment across clusters, there is also significant partitioning of genes at the functional level, with differential enrichment of Pfam annotated genes across clusters (Fig 5). Pfams are enriched by an average value of 0.25 and a standard deviation of 0.10, ranging from 0.13 in cluster 4 to 0.32 in cluster 2. This enrichment is significant (p<0.01) for most of the clusters (Fig 5c). This result suggests that clusters 1, 2, and 4 have significant functional enrichment, while functional enrichment on cluster 3 is marginally significant. Genes from a particular Pfam are most often assigned to only two or three clusters (Fig 5b). While functional enrichment in general is less strong than taxonomic enrichment, this may be due to the relative coarseness of functional annotation compared to taxonomic assignments, and our inability to annotate many genes with confidence.

**Fig. 5.**
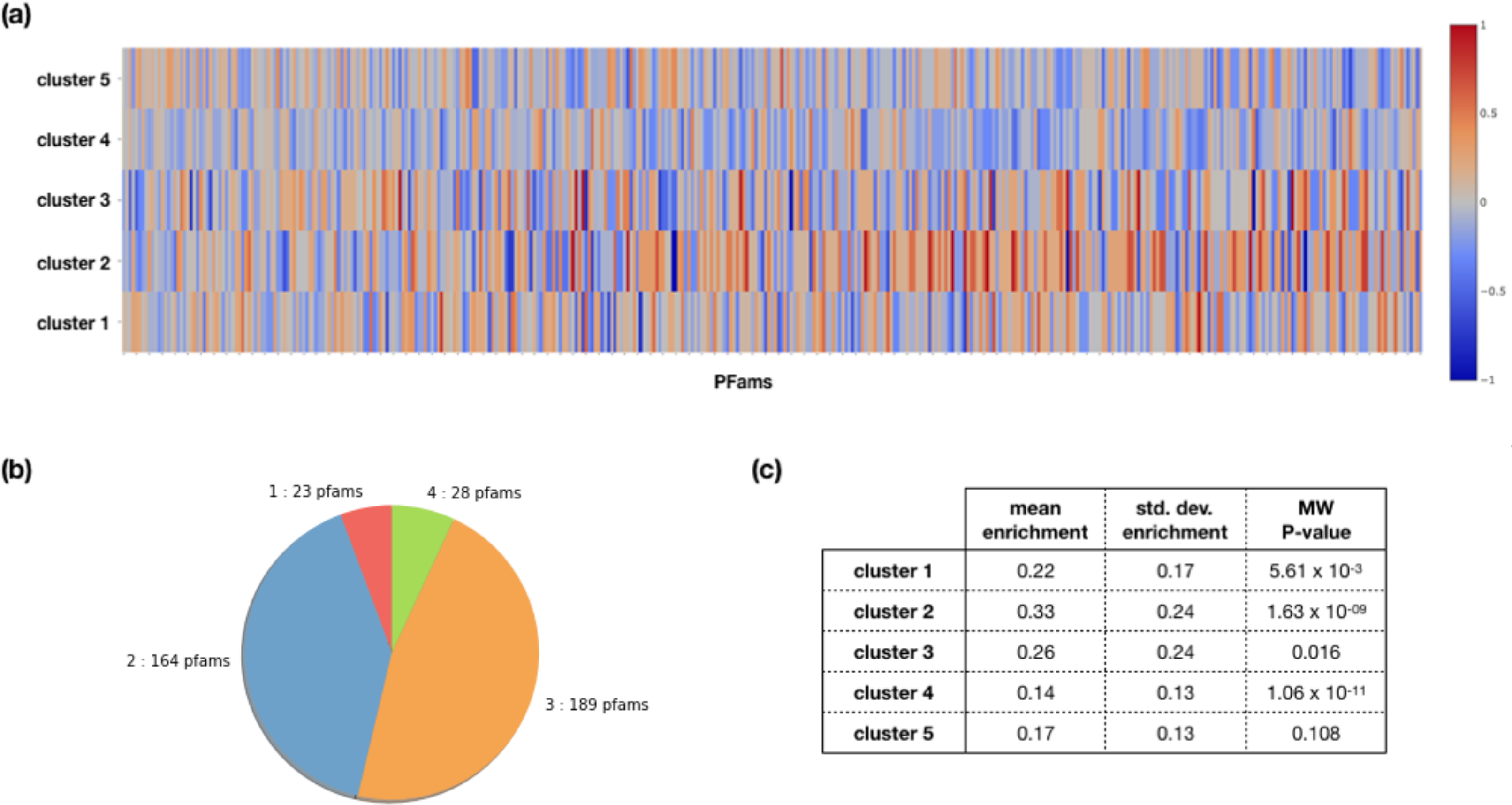
Functional enrichment in clusters. (a) Heatmap of enrichment (red) or depletion (blue) of the 405 most abundant Pfam families (columns) across each cluster (rows). Pfams are ordered left to right by the number of genes annotated to it, from the most abundant Pfams to the Pfams with as few as 20 genes annotated to it. (b) Pie chart of the number of clusters in which Pfams are enriched. (c) Mean positive enrichment value, standard deviation of positive enrichment values, and the Mann-Whitney P value for significance of cluster enrichment, for each cluster.

Enrichment of specific Pfams corresponding to some ecologically important functions indicate possible differentiation in ecological function between clusters. For example, glycosyl hydrolase family 3 (Pfams PF00933, PF01915), which corresponds to exo-acting glucosidases, is enriched across clusters 3, 4, and 5, and depleted in clusters 1 and 2, while glycosyl hydrolase family 16 (Pfam PF00722), which corresponds to endo-acting glucanases, is enriched strongly on cluster 3, depleted in clusters 1 and 2, and near the null value for clusters 4 and 5 (SI Fig 8). Proteorhodopsin, a photoactive transmembrane proton pump first identified in bacteria in SAR86 [31] and used by SAR86 for photoheterotrophic ATP generation, is enriched in clusters 3 and 4, and depleted in clusters 1, 2, and 5 (SI Fig 9).

## Discussion

While SAR86 is generally considered to be a ubiquitous heterotroph in the ocean, this study demonstrates that SAR86 harbors immense within-species genetic diversity that is strongly associated with environmental variables. These distinct environmental distributions of gene clusters define a deeper geographic variability within the SAR86 clade than previously appreciated. The three near-complete and two partial genomes available for SAR86 [25, 26] show high diversity within this clade; average nucleotide identity between genomes is between 70-80% (SI Table 5). In light of this high diversity, it is perhaps not surprising that the geographically variable genes in the SAR86 pangenome can be decomposed into five distinct clusters with different geographic distributions associated with unique environmental variables. These clusters are differentiated at the taxonomic and functional level, which has implications for our understanding of the biogeography of SAR86, as well as its ecological role within microbial communities in the marine environment.

Using a data intensive approach to build machine learning models of the relationship between SAR86 genes and environmental variables at a global scale, we demonstrate how such an approach can be used to better understand the factors shaping the biogeography of microbial clades. This approach can reveal patterns that would likely be missed at the 16S OTU or community level, or using data from a smaller scale. Particularly as metagenomics data become increasingly available in the future, such an approach holds promise for illuminating the relationship between microbial community structure and ecological function across broad taxonomic and spatial scales.

The results of this study identify clusters of genes that, while their phylogenetic relatedness is unknown, are taxonomically and functionally differentiated and occupy distinct environmental distributions. While the functional traits that confer niche restriction within these distributions is not obvious from our results, functional differentiation across clusters of glycosyl hydrolases (SI Fig 8) – an important class of enzymes for heterotrophic metabolism of polysaccharides – and proteorhodopsin (SI Fig 9) – a light-driven means of energy generation and enhanced nutrient and organic carbon uptake – suggest that genes associated with different clusters define distinct functional roles filled by each cluster. Glycosyl hydrolase families 3 and 16 target many of the same substrates – β-linked glucans, including the abundant marine plankton storage glucan laminarin – but using different enzymatic mechanisms [32]. The strong enrichment in cluster 3, and strong depletion in clusters 1 and 2, of both families, compared to the enrichment of only family 16 in clusters 4 and 5, may indicate distinct ecological functions of SAR86 across clusters that utilize differing metabolic strategies and have disparate impacts on carbon remineralization. Proteorhodopsin genes are only enriched in clusters 3 and 4, the two clusters associated with lower latitudes and more abundant sunlight, and are depleted in clusters 1 and 5, which are associated with temperate latitudes. This latitudinal pattern may also indicate distinct energy generation and metabolic strategies that correspond with the environmental distributions of the clusters. Given the clear taxonomic and functional partitioning of the SAR86 pangenome across clusters with distinct geographic distributions associated with unique environmental conditions, we conclude that the clusters described here define previously unidentified ecotypes within the SAR86 clade.

The geographic distributions of SAR86 ecotypes are consistent with previous studies. An investigation of temporal and geographic patterns in SAR86 noted that while the phylogenetic substructure of the SAR86 clade implies that it may be made up of multiple ecotypes, these could not be identified at the limited geographic resolution of the study [24]. The potential existence of SAR86 ecotypes was also noted in the apparent geographic distributions of SAR86A, B, C, and D genomes [25], which differed in their distributions across coastal versus open ocean sampling sites and along temperature gradients. This general observation is supported by the predicted distributions of the clusters identified in our study, for which three clusters (clusters 2, 3, and 4) are partially defined by their warmer, open ocean distributions, and two (clusters 1 and 5) are associated with cooler temperatures. The difficulty of identifying ecotypes in SAR86 contrasts with SAR11, for which distinct ecotypes have been identified within a constrained geographic sample because they were strongly associated with differences in depth and salinity distributions [15]. This study was able to identify SAR86 ecotypes, despite their partially sympatric distributions that cause single sampling sites to be composed of genes from multiple clusters, because of the larger data size and geographic distribution of the TARA dataset, and our unique approach to defining ecotypes based on quantitative models of environmental associations with geographically variable genes. Whereas ecotypes are typically identified by building a phylogeny based on core genes and observing whether environmental variables map over the phylogeny [e.g. 23, 33], our approach is quantitative, objective and independent of a priori knowledge of phylogeny, and results in sets of genes and functional features that define the ecotype.

The taxonomic and functional differentiation of genes across SAR86 ecotype clusters is significant in the context of interactions between microbial community structure, function, and ecology. Both community composition [16–18, 34] and functional traits [3, 4, 8, 19] vary geographically and can be predicted to some extent by environmental variables [8, 17]. Taxonomic variation can lead to functional differentiation of microbial communities [4, 35, 36], which ultimately shapes biogeochemical cycling and ecosystem function; conversely, functional redundancy across microbial taxa can complicate the relationship between structure and function [37], with taxonomically variable communities playing similar functional roles [38]. Disentangling the relationship between environment, biogeography, structure, and function is therefore a significant ongoing challenge in microbial ecology [5, 7, 8, 10]. By focusing on patterns at the individual gene level within a single clade, we are able to uncover patterns in environmental distributions of genetic diversity at a scale that would normally be obscured by the complexity inherent to microbial communities. For example, previous studies have found that functional classifications of taxa are better predicted by environmental parameters than taxonomic 16S-based classifications [8]; however, these functional classifications are broad – all of the SAR86 pangenome would be classified as ‘aerobic chemoheterotroph’ – in order to control for the vast genetic diversity of traits in mixed microbial communities. It is likely that within the SAR86 pangenome there is ecological differentiation within this category that, for example, could lead closely related phylotypes of SAR86 that belong to different ecotypes to utilize different substrates [33, 39, 40]. This hypothesis is supported by the functional enrichment across our clusters and the differential enrichment of carbohydrate utilizing enzymes (SI Fig 8). Previous analyses of the genomic context of SAR86 genomes also suggest that much of the diversity among SAR86 genomes may be driven by fine scale diversification of catabolic enzymes on loci associated with TonB dependent receptors [25], which are responsible for transporting carbon compounds (as well as metals) into the cell [41].

The accuracies of our gene models are better on average than previous studies (0.79 vs 0.48, [8]), which may similarly be due in part to our focus on modeling individual genes rather than whole communities. This difference in model accuracy may also be due to our consideration of different, and a larger number, of input environmental features. Here, the environmental features were chosen for their availability at global resolution rather than their human-predicted importance in regulating microbial function. These environmental features may be more predictive of the distributions of SAR86 genes, even if they are less relevant to biological function. The environmental factors that influence whether an organism grows in a particular location or community may be different from those that drive their function within that community: for example, an organism may only grow in fresh or saline waters, while the maintenance of a nitrogen fixation pathway depends on nutrients or other factors. It is important to note that those environmental features that are selected as most predictive for each gene model do not necessarily drive the growth of SAR86 in a causal manner, but implies only that these environmental features are good predictive proxies for the presence of that gene. The interpretation of the most predictive environmental features may vary depending on the feature; some features may be a proxy for biological phenomena, while others simply define oceanographic regions, or are proxies for other factors that cannot be measured that are true causal drivers of variation. The features chosen by the L1 regularization procedure are also likely biased by the scope of the samples used as inputs to the model. For example, the cluster associated with western hemisphere longitudes is overrepresented in sites from the Pacific Ocean in the TARA expedition dataset. However, there are longitudes both east and west of the antemeridian in the Pacific, represented as negative and positive longitudes in the models, and it is a limitation of the TARA dataset that only samples from the eastern part of the basin, in the western hemisphere, are represented. This limitation results in an unnaturally sharp transition in cluster projections on the antemeridian in the Pacific Ocean for those clusters for which longitude is a strong predictor. This observation also serves as a note of caution for the interpretation of the global projections, whose predicted distributions will likely break down most in locations for which representation of samples is most sparse, e.g. in polar regions.

We are able to make accurate predictions of geographic distributions of SAR86 genes, identifying previously unknown biogeographical complexity within an otherwise ubiquitous heterotrophic clade and making global projections of the distributions of SAR86 ecotypes associated with distinct environmental distributions. Our modeling approach leverages a large dataset across broad geographic regions, demonstrating the potential of machine learning and the use of broader scale integrated datasets for marine microbial ecology. The five global ecotypes underlying the highly diverse SAR86 clade, the taxonomic and functional differentiation across ecotypes, and the distinct environmental distributions of SAR86 genetic diversity highlight the importance of SAR86 within marine microbial communities and broadens the context for interpreting their ecological impact across the world’s oceans.

## Acknowledgements

This work was supported by a Deep Carbon Observatory Deep Life Modeling & Visualization Fellowship to AH; OCE-1736772 to CA; Gordon and Betty Moore Foundation grant #3300 to KSP; and grants from the Beyster Family Fund of the San Diego Foundation and Life Technologies Foundation to JCVI.

## Conflict of Interest

The authors declare no conflict of interest.

## Author Contributions

CD and SY created the SAR86 co-assembly of SAR86 genes from the Global Ocean Sampling sequences, and CD annotated the SAR86 pangenome. SN created the pangenome and mapped TARA samples to the SAR86 pangenome. AH gathered satellite environmental data, created the models, did clustering, identified ecotypes and analyzed data. JL gathered historical environmental data. All authors contributed to discussion of data and writing of the manuscript.

