## Supplemental Information for "Global ecotypes in the ubiquitous marine clade SAR86"

**List of Content**

SI Figures 1-9

Captions for External SI Tables 1-6

Supplemental Text

1 – Supplemental Methods

2 – Supplemental Results

References for Supplemental Information

**Figures**

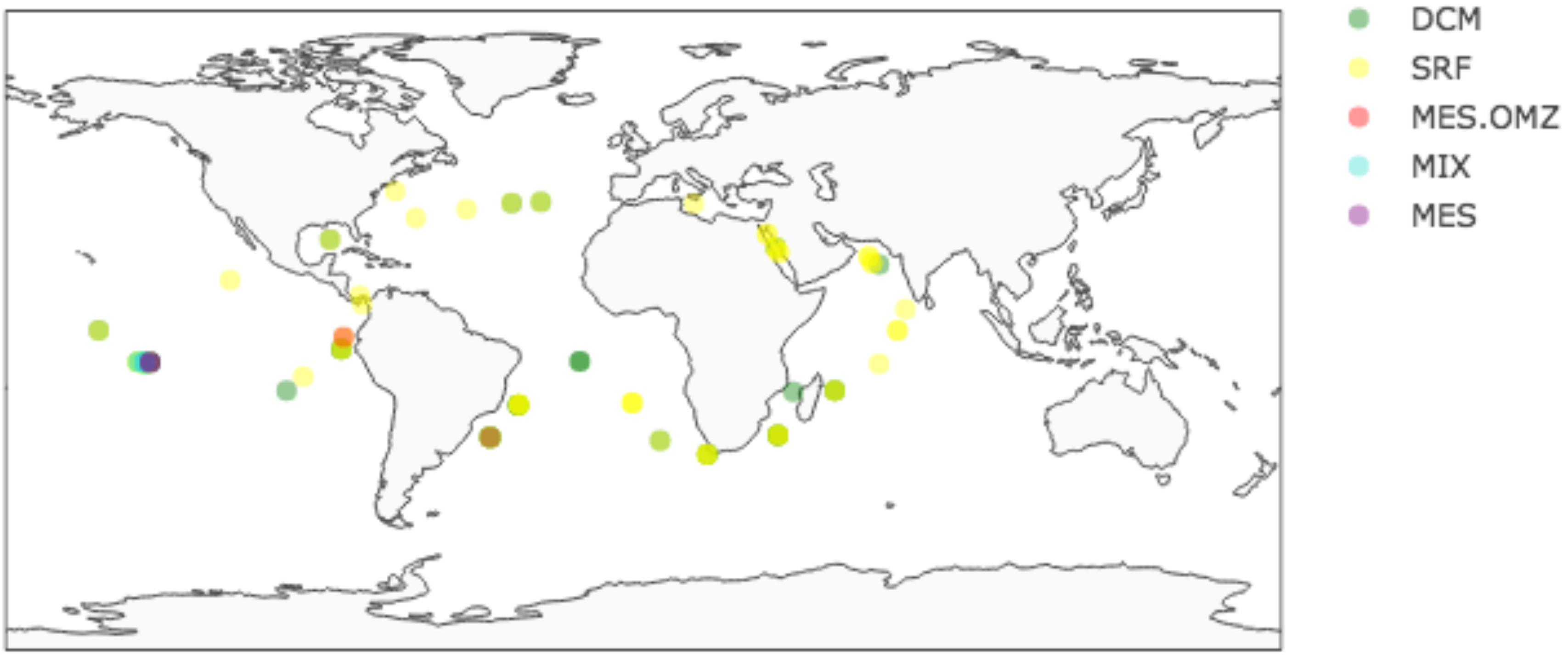


**SI Fig. 1** – Map of TARA station sampling locations. Points are colored by depth region sampled: DCM: deep chlorophyll maximum; SRF: surface; MES.OMZ: mesopelagic in ocean minimum zone; MIX: mixotrophic; and MES: mesopelagic.

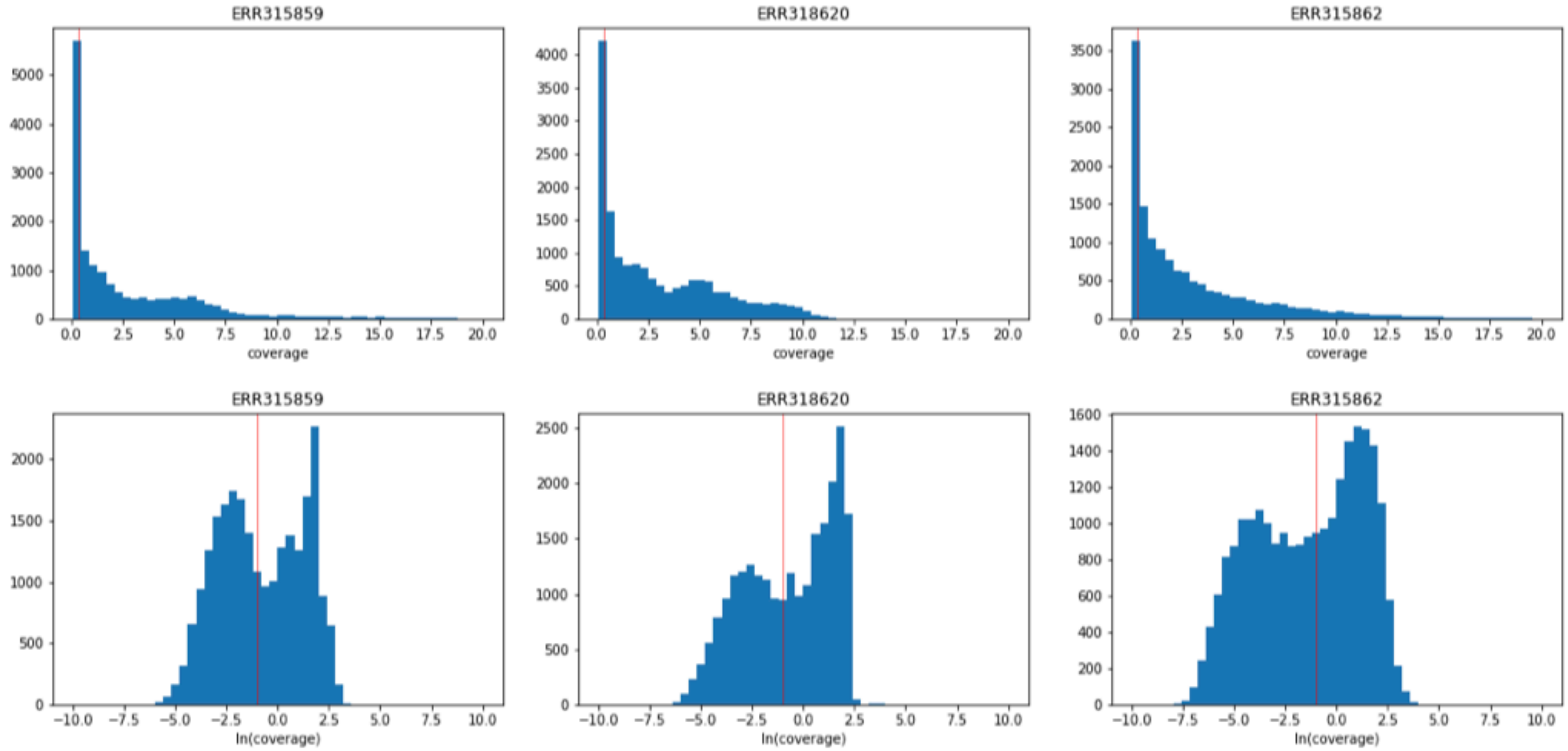


**SI Fig. 2** – Representative figures of the distribution of gene coverage (top row) and natural log of gene coverage (bottom row) for three example TARA sites ERR315859, ERR318620, and ERR315862. The coverage threshold that was used for presence/absence cutoffs is indicated by the vertical red line.

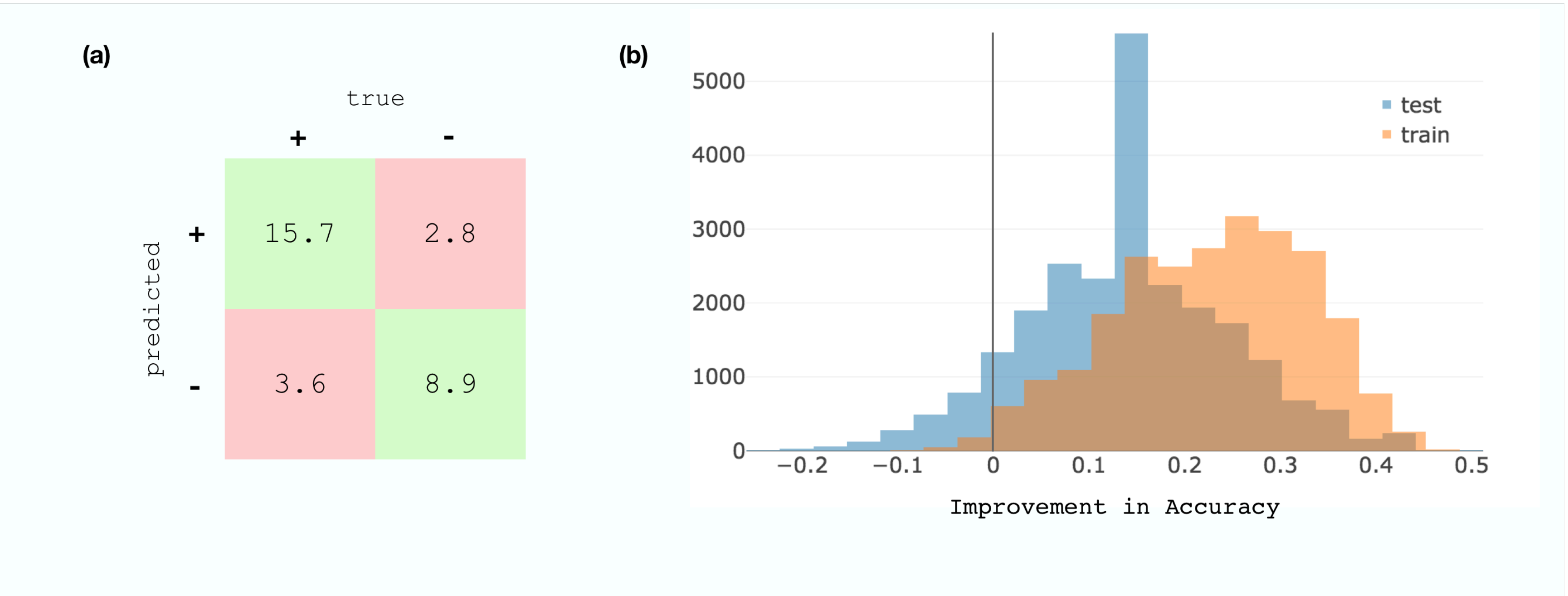


**SI Fig. 3** – (a) Average confusion matrix over all gene models. True labels are indicated by columns, while labels predicted by the models are indicated by rows. From top left to bottom right rowwise, this corresponds to the average number of true positive, false positive, false negative, and true negative predictions in the test set. (b) Histogram of improvement accuracy for the test set (blue) and training set (orange). Improvement accuracy is defined as improvement over the majority class accuracy (see text). An improvement accuracy great than zero, indicated by the vertical line, corresponds with models that score better accuracies than the majority class accuracy.

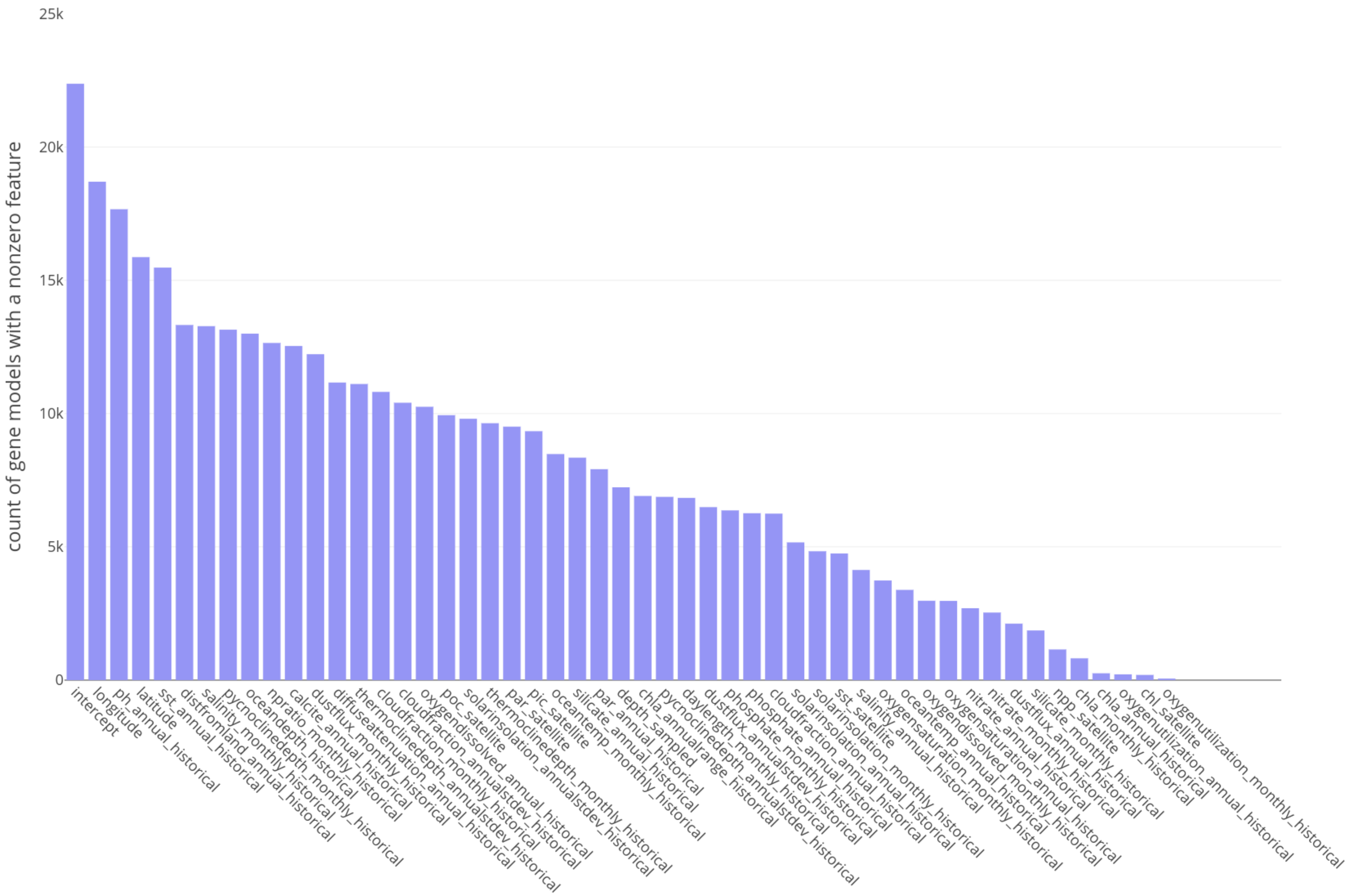


**SI Fig. 4** – Number of gene models (y axis) for which a particular environmental feature (x axis) was selected as nonzero during logistic regression model training. Features are sorted in order of the number of gene models for which it was nonzero.

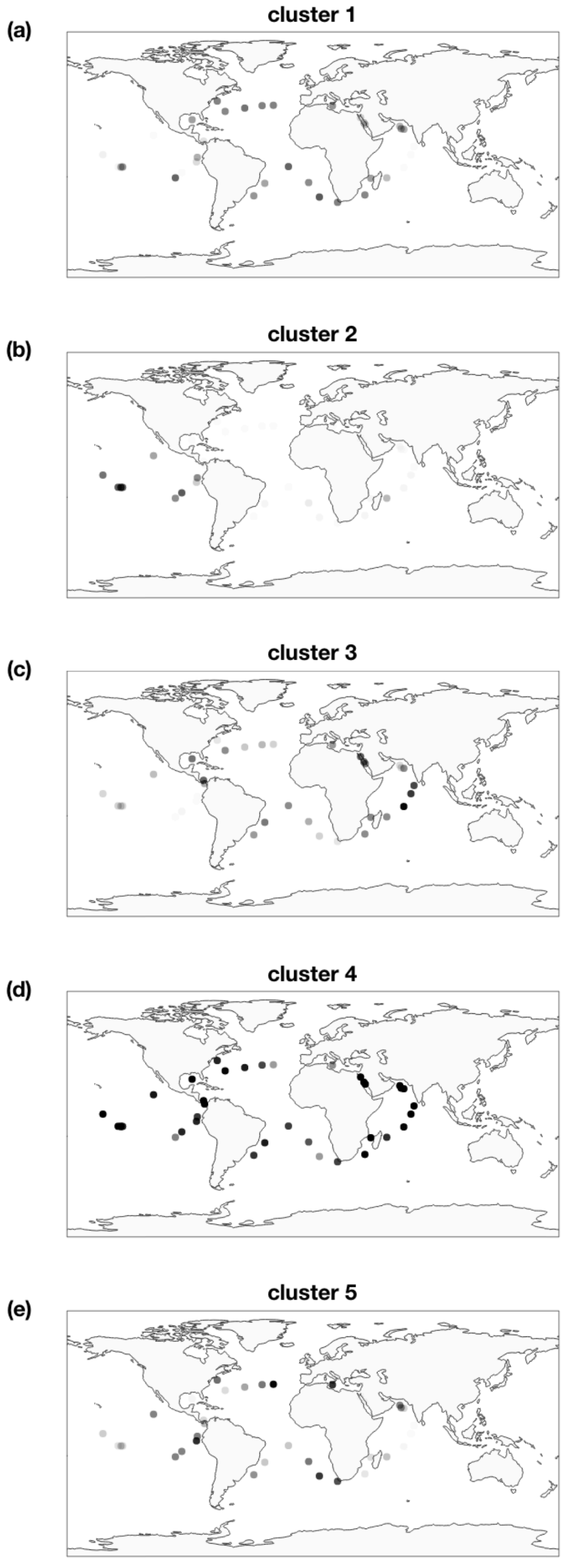


**SI Fig. 5** – Map of the relative proportion of geographic clusters 1-5, top to bottom, at each TARA site. The opacity of each symbol indicates the percentage of genes from that TARA site that belong to that cluster.

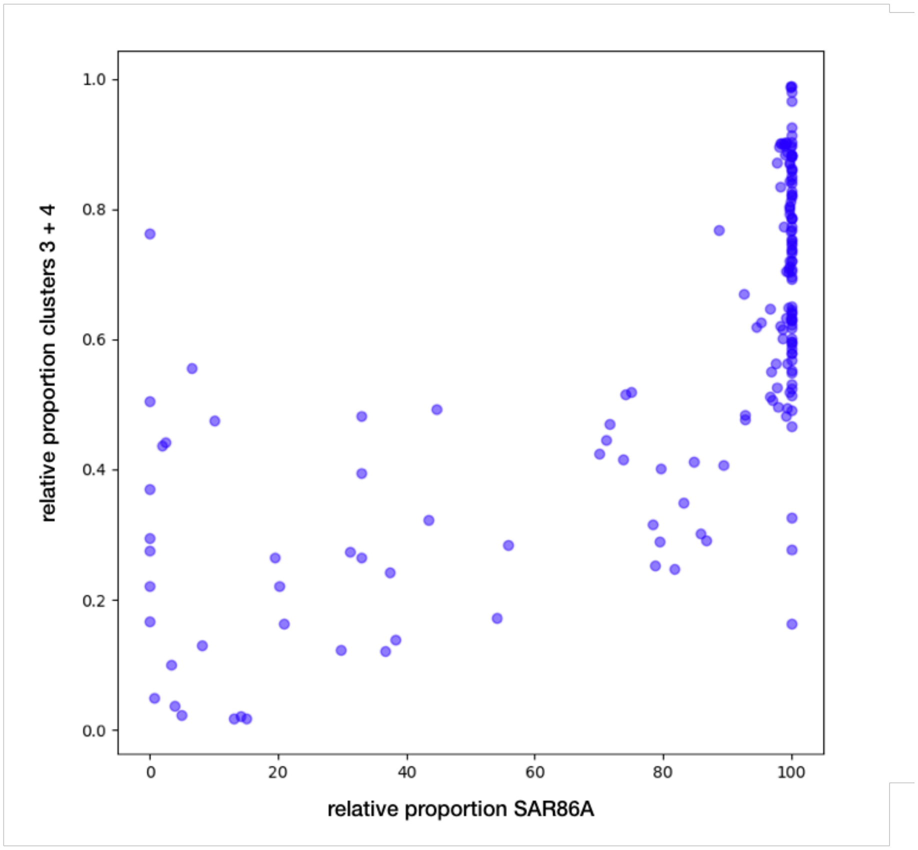


**SI Fig. 6** – Correlation between the relative abundance of SAR86A (vs SAR86E) and the relative proportion of clusters 3 + 4 (vs clusters 1 + 5) at each TARA site (blue dots). Pearson $R^2 = 0.70$, $P = 1.56\times10^{-26}$.

| | c1 | c2 | c3 | c4 | c5 |
|---|---|---|---|---|---|
| **c1** | 0.29 | | | | |
| **c2** | -0.11 | 0.69 | | | |
| **c3** | -0.06 | -0.18 | 0.53 | | |
| **c4** | -0.01 | -0.04 | 0.18 | 0.21 | |
| **c5** | 0.07 | -0.09 | -0.05 | -0.06 | 0.26 |

**SI Fig. 7** – The average correlation coefficient of normalized read coverage across TARA sites for genes from each cluster compared to every other cluster. c1=cluster 1, c2 = cluster 2, and so on.

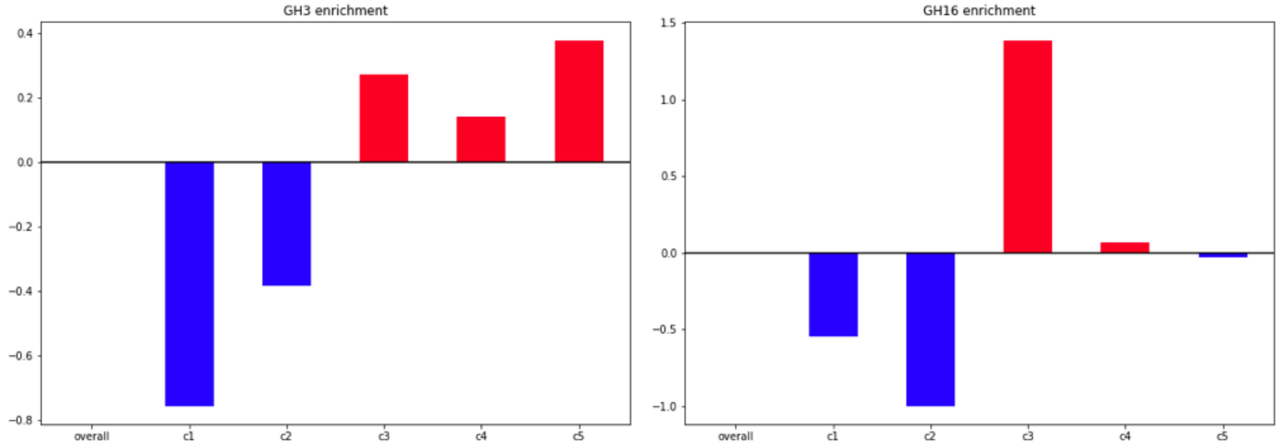


**SI Fig. 8** – Functional enrichment of Pfams associated with glycosyl hydrolase family 3 (left) and glycosyl hydrolase family 16 (right) in each cluster. The expected value ('overall') is indicated by a horizontal line at 0, enriched values (red) appear above this line and depletion values (blue) below this line.

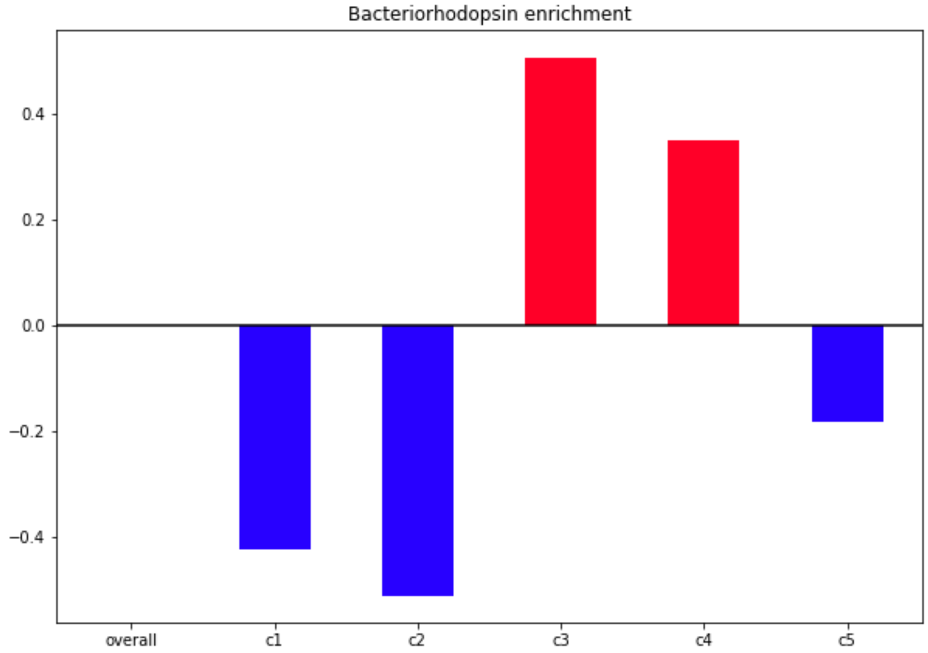


**SI Fig. 9** – Functional enrichment of Pfams associated with proteorhodopsin in each cluster. The expected value ('overall') is indicated by a horizontal line at 0, enriched values (red) appear above this line and depletion values (blue) below this line.

### External Table Captions

**SI Table 1** – Metadata for each of the 51 environmental variables input to the logistic regression gene models, including variable name, time span metric over which historical sources are averaged or sample/satellite sources are gathered, original units of the dataset, whether data is available a depth resolution or surface only, and the original data source.

**SI Table 2** – Satellite and historical environmental data corresponding to the sampling site, date, and depth of each TARA site.

**SI Table 3** – The cluster centroid coefficient for each environmental feature. Columns: 'feature', environmental feature name; cluster1-5: centroid coefficient for that cluster.

**SI Table 4** – Cluster membership proportions and Shannon Diversity metric for each TARA site. Columns: 'run_id', the European Nucleotide Archive run accession ID; 'longitude' and 'latitude': coordinates at which TARA site was sampled. 'tara_label': TARA project label. 'cluster1-5': proportion of genes present at that site assigned to that cluster. 'shannon': shannon diversity metric for each TARA site.

**SI Table 5** – Average nucleotide identity pairwise comparisons of SAR86A-E. Each cell is the ANI value for genome 1 (rows) compared against genome 2 (columns). ANI calculated after [1] using the online calculator at https://ani.jgi-psf.org/html/calc.php?.

**SI Table 6** – Assembly statistics for the massive co-assembly of GOS metagenomes.

### Supplemental Text

### 1 – Supplemental Methods

#### *1.1 Co-assembly generation*

The co-assembly of SAR86 sequences was generated from 226 metagenomic samples from the Global Ocean Sampling ([2], GOS) to create a *de novo* SAR86 database (Genbank Bioproject PRJNA13694 and European Bioinformatics Institute accession numbers ERX913362-ERX913706). Raw reads from all metagenomes sequenced on pyrosequencing (50 samples) and Sanger (226 samples) platforms were co-assembled using the CELERA assembler [3] at 92% nucleotide identity (SI Table 6). This threshold allowed for consensus assemblies at the species and strain level [4] with reasonable computation times. The resulting scaffolds encompass 3 Gbp of contiguous DNA sequence, and 85% of the sequence reads could be mapped back to the assembly. Open reading frames (ORFs) on scaffolds were called using MetaGene [5]. To determine the putative phylogenetic origin of the scaffolds, each predicted peptide was phylogenetically annotated using Automated Phylogenetic Inference System (APIS) [6], which annotates according to the position of the peptide within a phylogenetic tree. Thus, a peptide 99% similar to a SAR86 protein will be annotated as SAR86 (with the associated taxonomic tree), while a peptide that branches basally within the phylogenetic tree next to *Gammaproteobacteria* would only be annotated as such. The scaffolds were taxonomically annotated at the lowest level for which greater than 50% of the ORFs had agreement in the APIS calls. This approach has been used in previously published pangenome biogeographic analysis [7]. Scaffolds assigned to SAR86 were used in the metagenomic SAR86 co-assembly that was fed into the MIDAS tool for SAR86 pangenome creation.

#### *1.2 Pangenome creation*

A custom pangenome of SAR86 genes was created from 5 partial or near-complete reference genomes (SAR86A-E, [8, 9]), 4 single-cell sequencing draft genomes (PATRIC ID 1007118.3, 1007119.3, 1009425.3, and 1009426.3, [10]), and SAR86 genes from the massive co-assembly of GOS metagenomes (Supplemental Text 1.1). The SAR86 pangenome was created with the MIDAS tool [11]. 58 423 genes were clustered at 90% nucleotide identity, resulting in a total of 51 711 nonredundant SAR86 genes in the SAR86 pangenome. Only 4 188 (8%) of these genes were derived from the five reference SAR86 genomes, highlighting the diversity of SAR86 genes that are still without genomic reference.

#### *1.3 Global SAR86 gene presence/absence details*

A global dataset of SAR86 gene presence/absence was created by mapping sequencing reads from globally-distributed shotgun metagenomes from the TARA project [12] to the SAR86 pangenome. TARA was a large-scale, multi-year effort to sequence hundreds of marine seawater samples from globally-distributed sites, and from depths ranging from surface to mesopelagic, using consistent techniques (SI Fig 1). To date, the TARA project has made available 243 individual metagenomic samples, 198 of which were for live samples (pore size >0.22) and 45 of which were <0.22 filter pore size blank controls. MIDAS (v1.1.0, [11]) was used to perform read alignment of the 198 live metagenomes with bowtie2 (options --very-sensitive) and using the best hit with a % DNA identity >90% for each read. Reads with mapping quality <20 or mean base quality <20 were discarded. Where multiple sequencing runs had been conducted for a single sample, the run with the highest number of sequencing reads was used.

This mapping procedure resulted in gene coverage values for each of the 51 711 SAR86 genes at each of the 198 live TARA sites. Gene coverage was normalized by dividing observed coverage by the mean coverage at that site, so that each site had a mean normalized coverage value of one. This continuous value was converted to a binary gene presence/absence at a threshold of 0.37, where genes with a normalized coverage value less than or equal to 0.37 were considered absent from the sample, and those with coverage greater than 0.37 were considered present. This threshold value was chosen from manual inspection of the SAR86 gene coverage distribution at individual TARA sites; where a strain or mixture of strains of SAR86 are present, a peak in normalized gene coverage frequency is observable, while erroneous gene assignments form a long tail of low coverage. The gene coverage values of this erroneous long tail was typically below $e^{-1}$, or 0.37, as was particularly visible when viewed as a natural log distribution of gene coverage (SI Fig 2). This value of 0.37 was thus chosen as a threshold below which normalized SAR86 genes are 'absent'.

#### *1.4 Environmental data curation details*

51 environmental features from contemporary satellite data and historical averages were curated for input to the logistic regression models (Methods, Supplemental Text 1.5) and for global projections of clusters (Methods, Supplemental Text 1.6). These features, their sources, definitions, and units are summarized in SI Table 1. These data included 6 features from contemporary satellite data – sea surface temperature, chlorophyll *a* concentration, photosynthetically active radiation at the sea surface, particulate inorganic carbon at the sea surface, particulate organic carbon at the sea surface, and net primary productivity at the sea surface. Three features constituted data about the location of the sample: latitude, longitude, and

depth sampled. The remaining features were environmental data available from historical averages of many values of interest, such as day length, dust flux, pycnocline depth, apparent oxygen utilization and concentrations of nitrate, phosphate, and silicate (SI Table 1). Most historical data represented the 57-year average annual or monthly value for that feature. Other historical data features represented decadal average values for that feature. For example, the historical annual mean pH value, derived from BioOracle [13], is the average annual pH over the period 1955-2012; the historical monthly mean nitrogen:phosphate ratio, in contrast, is the average N:P ratio during a particular month, that occurred over the period of 1955-2012. The annual standard deviation for a historical value, available for some historical environmental features, provides information about the typical variation in an environmental variable over the course of a year for the relevant historical period.

For each TARA site, the environmental feature value closest to the sampling site's latitude, longitude, and, where relevant, the sampling depth and/or sampling date, was derived from the original environmental data source. Environmental features, which are all continuous variables, were then centered and normalized to a mean of zero and a standard deviation of one across all TARA sites. This preprocessed environmental feature matrix (SI Table 2) served as the input feature vectors for each TARA site during model training.

***1.5 Model training and parameter tuning***

Environmental feature vectors for only those 155 TARA sites where SAR86 was present and no environmental data were missing were used as inputs for model training. First, we dropped 22 of the 198 TARA sites where SAR86 was not present or abundance was low (defined as fewer than 1000 SAR86 genes with at least 5x coverage). 20 of these 22 sites were mesopelagic; however, 41 sites from mesopelagic depths had SAR86 abundances sufficiently high to be included in model training. An additional 21 sites for which environmental data were missing were also filtered out. The resulting 155 TARA sites were randomly split into training, validation, and test sets of 111, 13, and 31 sites respectively. The training set was used for creating the logistic regression models for each gene; the validation set was used for parameter tuning of the regularization penalty parameter; and the test set was used to evaluate the final trained model performance. Validation and test sets using samples independent of the training set provide a more accurate measure of model performance than training set accuracy, which may provide an overly optimistic measure accuracy due to overfitting. These set sizes were chosen to maximize our relatively small amount of training data while still maintaining sufficient data in the validation and test sets to obtain useful parameter tuning and model accuracy statistics. The 20 TARA samples that were not used in model training, for which SAR86 abundance was low and environmental data was complete, were used to test the accuracy of the models in sites outside the distribution of training data. 18 of these 20 sites were from mesopelagic samples, and originated from Indian, Atlantic, and Pacific ocean basins.

Of the 51 711 genes in the SAR86 pangenome, 24 317 were present at 20-80% of TARA sites. Logistic regression models were trained only for these variable genes, since it was not meaningful to predict the distribution of either very rare or very common genes.

L1 regularization was used during model training to select the most predictive subset of environmental features, while the coefficients for less predictive environmental features converge to zero so that these features are effectively ignored. This approach enables us to identify which of the many environmental variables are most reliably associated with a gene's presence, while reducing overfitting. The L1 regularization strength was tuned to minimize overfitting while maximizing accuracy in the validation set. Penalty parameters ranging from 0 (strongest regularization) to 1 (no regularization) were tested, and model performance in the validation set was used to evaluate the penalty parameter. A penalty parameter of 0.7 was chosen, which improved overfitting (66.9% of models were overfit as compared to 70.1% of models with no regularization penalty) but achieved comparable mean validation accuracy (82.1% across all models, as compared with 82.2% accuracy in the validation set with no regularization penalty). At this regularization strength, an average of 17 out of 51 environmental features were selected during model training as the most predictive environmental features (i.e. were associated with nonzero model coefficients).

The recall, or the sensitivity rate, was calculated as (TP/(TP + FN); precision is defined as (TP/(TP + FP); and the F1-score is defined as 2*((precision*recall)/(precision+recall)), where TP = true positive; FP = false positive; FN = false negative.

#### *1.6 Clustering and cluster projection details*

SAR86 genes were clustered into five clusters based on the similarity of the environmental feature coefficients derived during model training (Methods, Supplemental Text 1.5). Clustering on environmental feature vectors was advantageous over identifying clusters based on, for example, gene covariation across TARA sites, which would identify genes with similar geographic distributions but would not uncover associations between genes and environmental conditions nor would it be possible to extrapolate gene distributions beyond the TARA sampling sites.

The number of clusters, $k=5$, was chosen as the value of $k$ at which the inertia (sum of the squared distance of each point to its centroid) begins to decrease less rapidly; where the distance between centroids begins to increase less rapidly; and where map projections and comparisons of Euclidean distances of cluster centroids look unique for each cluster without producing repetitive distributions. Exploratory analysis in a Jupyter notebook and a python script for reproducing clusters are available on the Github repository [14].

Using the cluster centroid coefficients for environmental features associated with each cluster, global projections of predicted cluster presence were created. Predictions of cluster distributions were derived by multiplying normalized environmental feature values for each latitude and longitude coordinate at global, 9km resolution by the cluster centroid coefficient for each environmental feature, summing this vector of coefficient-multiplied environmental feature values, and converting to a [0,1] scale using a sigmoid function. From the resultant prediction, any value greater than or equal to 0.5 is considered a 'present' prediction, anything less than 0.5 is considered 'absent', and how close to the extremes of 0 or 1 the prediction is gives a measure of the certainty of the prediction, with 0.5 the most uncertain prediction, 0 a very confident 'absent' prediction, and 1 a very confident 'present' prediction.

#### *1.7 Taxonomic & functional enrichment analysis details*

Of the 24 317 genes that were modeled, only 810 originated from one of the five reference SAR86 genomes; of these, 622 genes were from SAR86A, and 157 genes originated from SAR86E. Only the genomes SAR86A and SAR86E were thus used for genome enrichment analysis because only 5, 7, and 19 genes originated from SAR86C, D, and B respectively.

Genome distribution across clusters was evaluated by counting the percentage of genes in the SAR86 pangenome originating from SAR86A or SAR86E assigned to each cluster. The correlation of SAR86A/E relative abundance with their associated cluster proportion within

TARA sites was evaluated by deriving the relative abundance of SAR86A and E from the mapped TARA samples, and renormalizing such that the abundances of SAR86A and E sum to 1. The cluster proportions of cluster 1+5 and cluster 3+4 were similarly renormalized to sum to 1.

Enrichment values were calculated the same way for both contig and functional enrichment analyses. The enrichment of a particular contig or Pfam family annotation was calculated as (actual – expected)/expected, where the expected value is based on evenly proportioning cluster labels across contigs, and the actual value is the actual number of genes from a contig/Pfam assigned to the cluster. A contig/Pfam is enriched in a cluster if this statistic exceeds zero, with a value of e.g. 2.1 resulting when 210% more genes from that contig were assigned to a cluster than expected. A value below zero is a depletion relative to the expected value, and a minimum value of -1 is found where zero genes were assigned to a cluster. For each cluster, the enrichment vector of enrichment statistics for each contig/Pfam can then be used to test the null hypothesis that there is no difference in function or contig composition across clusters. To test the statistical significance of cluster enrichment vectors, a nonparametric Mann-Whitney U test was applied to test whether the vector of enrichment values associated with each contig/annotation for each cluster was significantly different from the expected distribution of all-zeros.

In the case of the functional enrichment analysis, there were 1337 Pfam families to which at least one gene was annotated. However, only 405 Pfams for which more than 20 genes were annotated to it were used for the enrichment analysis, since a rare Pfam with only one gene annotated to it will look perfectly enriched in whichever cluster to which that gene is assigned.

A Shannon diversity metric was used to measure the relative proportions of the five clusters at each TARA site. The Shannon metric accounts for both the relative evenness of the cluster proportions as well as how many of the five clusters are present at each TARA site. The minimum possible value of 0 would indicate a site at which genes from only one cluster was present, while the maximum possible value of $ln(5)$= 1.609 would indicate a site where genes from all five clusters are present in equal proportions. The Shannon diversity metric ranged from 0.699 to 1.532 across TARA sites (SI Table 4).

### 2 – Supplemental Results

#### *2.1 Cluster characteristics: highest magnitude coefficients*

The sign and magnitude of coefficients associated with each environmental variable is unique to each cluster (SI Table 3). Which environmental variables have the highest magnitude coefficients drives the predicted geographic distribution of each cluster (Fig 3).

In cluster 1, the environmental features with the highest magnitude coefficients included monthly historical ocean temperature (-1.23), monthly historical dust flux (-0.75), annual historical phosphate (-0.66), monthly historical pycnocline depth (0.50), and annual historical silicate (-0.40). The high magnitude of monthly features suggests a seasonal component, with genes more likely to be present during the winter and spring, while the negative coefficients associated with nutrients, temperature, and dust flux confine projections primarily to open ocean regions.

In cluster 2, the environmental features with the highest magnitude coefficients included longitude (-1.01), annual historical photosynthetically active radiation (PAR, 0.84), monthly historical cloud fraction (-0.72), and annual historical standard deviation of solar insolation (-

0.40). The PAR, cloud fraction, and solar insolation coefficients all select for higher likelihood of gene presence in areas of high light, few clouds year-round, and low variability in sunlight, as is typically found in lower latitudes, while the longitude coefficient selects for locations in the western hemisphere. The TARA samples from the Pacific Ocean were only sampled from the western hemisphere, and were also the only locations where mesopelagic samples were taken. Not coincidentally, the TARA sites with the highest proportion of cluster 2 genes present were the mesopelagic samples and those samples taken in the Pacific Ocean (Fig 4).

Cluster 3's most predictive environmental features included annual historical diffuse attenuation (-2.43), longitude (0.92), annual historical sea surface temperature (0.87), and monthly historical dust flux (-0.40). The global projections for cluster 3 predict gene presence most confidently in the eastern hemisphere, at lower latitudes where temperature is higher, and away from coasts where diffuse attenuation is lower.

Cluster 4 environmental features with highest magnitude coefficients included annual historical sea surface temperature (0.95), monthly historical dust flux (-0.53), longitude (0.42), annual historical pH (-0.41), and annual historical standard deviation in thermocline depth (-0.37). Similarly to cluster 3, the genes in cluster 4 are predicted more likely to be present in the eastern hemisphere and where temperature is higher. There is also a slightly higher likelihood of a gene being present in the southern hemisphere, where dust flux is lower. The coefficients of features associated with cluster 4 are more even than other clusters, and so additional features with lower coefficient magnitudes, such as a negative association with monthly historical phosphate (-0.26) and a positive association with annual historical standard deviation of cloud fraction (0.22) also help to explain the predicted distribution of cluster 4 genes.

Cluster 5 environmental features with highest magnitude coefficients included annual historical standard deviation in solar insolation (0.58), annual historical sea surface temperature (-0.49), contemporary particulate inorganic carbon (-0.42), annual historical cloud fraction (0.41), and annual historical silicate (-0.36). The positive association with variable solar insolation and negative association with temperature result in distributions of genes present at higher latitudes. This high-latitude distribution is also consistent with a positive association with cloud fraction, which is generally concentrated at higher latitudes and at the equator. The negative association with silicate restricts the southern polar distribution such that genes are less likely to be present near the Antarctic Convergence/AA Polar Front.
